# Genome-scale phylogeny and contrasting modes of genome evolution in the fungal phylum Ascomycota

**DOI:** 10.1101/2020.05.11.088658

**Authors:** Xing-Xing Shen, Jacob L. Steenwyk, Abigail L. LaBella, Dana A. Opulente, Xiaofan Zhou, Jacek Kominek, Yuanning Li, Marizeth Groenewald, Chris Todd Hittinger, Antonis Rokas

## Abstract

Ascomycota, the largest and best-studied phylum of fungi, contains three subphyla: Saccharomycotina (budding yeasts), Pezizomycotina (filamentous fungi), and Taphrinomycotina (fission yeasts); organisms from all three subphyla have been invaluable as models in diverse fields (e.g., biotechnology, cell biology, genetics, and medicine). Despite its importance, we still lack a comprehensive genome-scale phylogeny or understanding of the similarities and differences in the mode of genome evolution within this phylum. To address these gaps, we examined 1,107 genomes from Saccharomycotina (332), Pezizomycotina (761), and Taphrinomycotina (14) species to infer the Ascomycota phylogeny, estimate its timetree, and examine the evolution of key genomic properties. We inferred a robust genome-wide phylogeny that resolves several contentious relationships and estimated that the Ascomycota last common ancestor likely originated in the Ediacaran (~563 ± 68 million years ago). Comparisons of genomic properties revealed that Saccharomycotina and Pezizomycotina, the two taxon-rich subphyla, differed greatly in their genome properties. Saccharomycotina typically have smaller genomes, lower GC contents, lower numbers of genes, and higher rates of molecular sequence evolution compared to Pezizomycotina. Ancestral state reconstruction showed that the genome properties of the Saccharomycotina and Pezizomycotina last common ancestors were very similar, enabling inference of the direction of evolutionary change. For example, we found that a lineage-specific acceleration led to a 1.6-fold higher evolutionary rate in Saccharomycotina, whereas the 10% difference in GC content between Saccharomycotina and Pezizomycotina genomes stems from a trend toward AT bases within budding yeasts and toward GC bases within filamentous fungi. These results provide a robust evolutionary framework for understanding the diversification of the largest fungal phylum.

## Main

The fungal phylum Ascomycota is one of most diverse phyla of eukaryotes with ~65,000 known species that represent approximately three quarters of all known species of fungi^1^. The Ascomycota is divided in three subphyla. The Saccharomycotina subphylum is a lineage of more than 1,000 known species and 12 major clades^2^; commonly referred to as budding yeasts. Species in this lineage include the model organism *Saccharomyces cerevisiae*^3^ and several notable pathogens, such as the human commensal *Candida albicans^4^* and the multidrug-resistant emerging pathogen *Candida auris*^5^. The Pezizomycotina subphylum contains more than 63,000 described species in 13 classes^6^; commonly referred to as filamentous fungi. This subphylum contains several major plant and animal pathogens belonging to diverse genera, such as *Fusarium, Aspergillus, Zymoseptoria,* and *Magnaporthe*^71^. Finally, the Taphrinomycotina subphylum contains ~140 described species in 5 classes^6^; commonly referred to as fission yeasts. This subphylum contains the model organism *Schizosaccharomyces pombe* and the human pathogen *Pneumocystis jirovecii*^11,12^.

To better understand the evolution of species diversity and ecological lifestyles in Ascomycota fungi, a robust framework of phylogenetic relationships and divergence time estimates is essential. In the last two decades, several studies have aimed to infer the Ascomycota phylogeny, either using a handful of gene markers from hundreds of taxa^13–17^ or using hundreds of gene markers from tens of taxa^18–31^. To date, the most comprehensive “few-markers-from-many-taxa” phylogeny used a 6-gene, 420-taxon (8 Taphrinomycotina, 16 Saccharomycotina, and 396 Pezizomycotina) data matrix^13^, whereas the most comprehensive genome-scale phylogeny used an 238-gene, 496-taxon (12 Taphrinomycotina, 76 Saccharomycotina, and 408 Pezizomycotina) data matrix^22^ but was inferred using FastTree, a program that is faster but typically yields phylogenies that have much lower likelihood scores than those obtained by IQ-TREE and RAxML/RAxML-NG^23^. Key relationships supported by these studies include the monophyly of each subphylum and class and the sister group relationship of subphyla Saccharomycotina and Pezizomycotina. In contrast, relationships among classes are contentious between studies, particularly with respect to relationships between the 13 classes in Pezizomycotina^6^. For example, there is disagreement whether the sister group to the rest of classes in the Pezizomycotina is class Pezizomycetes^14^, class Orbiliomycetes^17^, or a clade comprised of both^19^.

Previous molecular clock-based estimates of divergence times for Ascomycota have all been based on few-markers-from-many-taxa data matrices^14,15,24–26^, resulting in age estimates for key events in Ascomycota evolution that have wide intervals. For example, analysis of a 6- gene, 121-taxon (1 Saccharomycotina, 118 Pezizomycotina, and 2 Taphrinomycotina) data matrix inferred that the origin of the phylum Ascomycota took place 531 million years ago (mya) (95% credibility interval (CI): 671-410 mya) (see their Scenario 4 in Table 3)^15^, while analysis of a 4-gene, 145-taxon (12 Saccharomycotina, 129 Pezizomycotina, and 4 Taphrinomycotina) data matrix inferred that the phylum originated 588 mya (95% CI: 773-487 mya)^14^. More importantly, the sparser taxon sampling of previous studies has prevented estimation of divergence times of several key divergence events of higher taxonomic ranks^24–26^ and stymied our understanding of their evolutionary pace. While these studies have significantly advanced our understanding of Ascomycota evolution, a comprehensive, genome-scale phylogeny and timetree stemming from the sampling of hundreds of genes from thousands of taxa from the phylum are still lacking.

A robust phylogenomic framework would also facilitate comparisons of genome evolution across the subphyla of Ascomycota. For example, the three subphyla differ in their genome sizes, with the genomes of Pezizomycotina species being notably larger (~42 Mb) than those of Saccharomycotina (~13 Mb) and Taphrinomycotina (~14 Mb)^27^. While several recent studies have analyzed major lineages within the two taxon-rich subphyla, Saccharomycotina^2,20,28^ and Pezizomycotina^29–31^, comparisons of genome evolution across the two subphyla are lacking. For example, a recent analysis of the tempo and mode of genome evolution in 332 Saccharomycotina found evidence of high evolutionary rates and reductive evolution across this subphylum^2^, but whether budding yeasts are faster evolving than filamentous fungi remains unknown. However, a recent analysis of 71 Ascomycota genomes showed that Pezizomycotina have much higher levels of gene order divergence than Saccharomycotina^21^. Similarly, genome-wide examinations of horizontal gene transfer events in dozens to more than a hundred Ascomycota genomes have revealed that Pezizomycotina acquired significantly higher numbers of genes from prokaryotic donors than Saccharomycotina ^32,33^. Although these studies have contributed to our understanding of certain evolutionary processes in the phylum, we still know relatively little about the evolution of Ascomycota genomes and their properties.

There are currently more than one thousand genomes from Ascomycota species that are publicly available, which span the diversity of Saccharomycotina (332 genomes representing all 12 major clades), Pezizomycotina (761 genomes representing 9 / 13 classes), and Taphrinomycotina (14 genomes representing 4 / 5 classes) (1,107 genomes as of December 14, 2018). These 1,107 genomes represent a much larger and representative source of genomic data across the entire Ascomycota phylum than previously available, providing a unique opportunity to infer a genome-scale phylogeny and timetree for the entire subphylum and compare the mode of genome evolution across its subphyla.

## Results and Discussion

### A genome-scale phylogeny of the fungal phylum Ascomycota

To infer a genome-scale phylogeny of Ascomycota fungi, we employed 1,107 publicly available genomes from species belonging to Ascomycota (Saccharomycotina: 332; Pezizomycotina: 761; Taphrinomycotina: 14) and six outgroups from the sister fungal phylum Basidiomycota. All genomes were retrieved from the NCBI GenBank database, ensuring that only one genome per species was included (Supplementary Tables 1 and 2). Analysis of genome assembly completeness reveals that 1,021/1,113 (~92%) genomes have more than 90% of the 1,315 full-length BUSCO genes^34^ (Supplementary Fig. 1).

1,315 BUSCO genes from 1,107 Ascomycota fungi and six outgroups were used to construct a phylogenomic data matrix (see Methods). After constructing the multiple amino acid sequence alignment and trimming ambiguous regions for each of these 1,315 BUSCO genes, we kept only the 815 BUSCO genes that had taxon occupancy of ≥ 50% for each subphylum (i.e., ≥ 7 Taphrinomycotina, ≥ 166 Saccharomycotina, and ≥ 381 Pezizomycotina) and whose amino acid sequence alignments were ≥ 300 sites in length. In the final set of 815 BUSCO genes, alignment lengths range from 300 to 4,585 amino acid sites (average = 690) and numbers of taxa range from 851 to 1,098 (average = 1,051) (Supplementary Table 3). The final data matrix contains 1,107 taxa, 815 genes, and 562,376 amino acid sites.

Inference using concatenation- and coalescent-based approaches yielded a robust, comprehensive phylogeny of the Ascomycota phylum (Fig. 1). The vast majority of internodes in both the concatenation-based (1,103 / 1,110; 99%) and the coalescent-based phylogeny (1,076 / 1,110; 97%) received strong (≥ 95%) support and were congruent between the phylogenies inferred using the two approaches; only 46 / 1,110 (4%) internodes were incongruent between the two phylogenies (Supplementary Figs. 2 and 3).

**Fig. 1.**
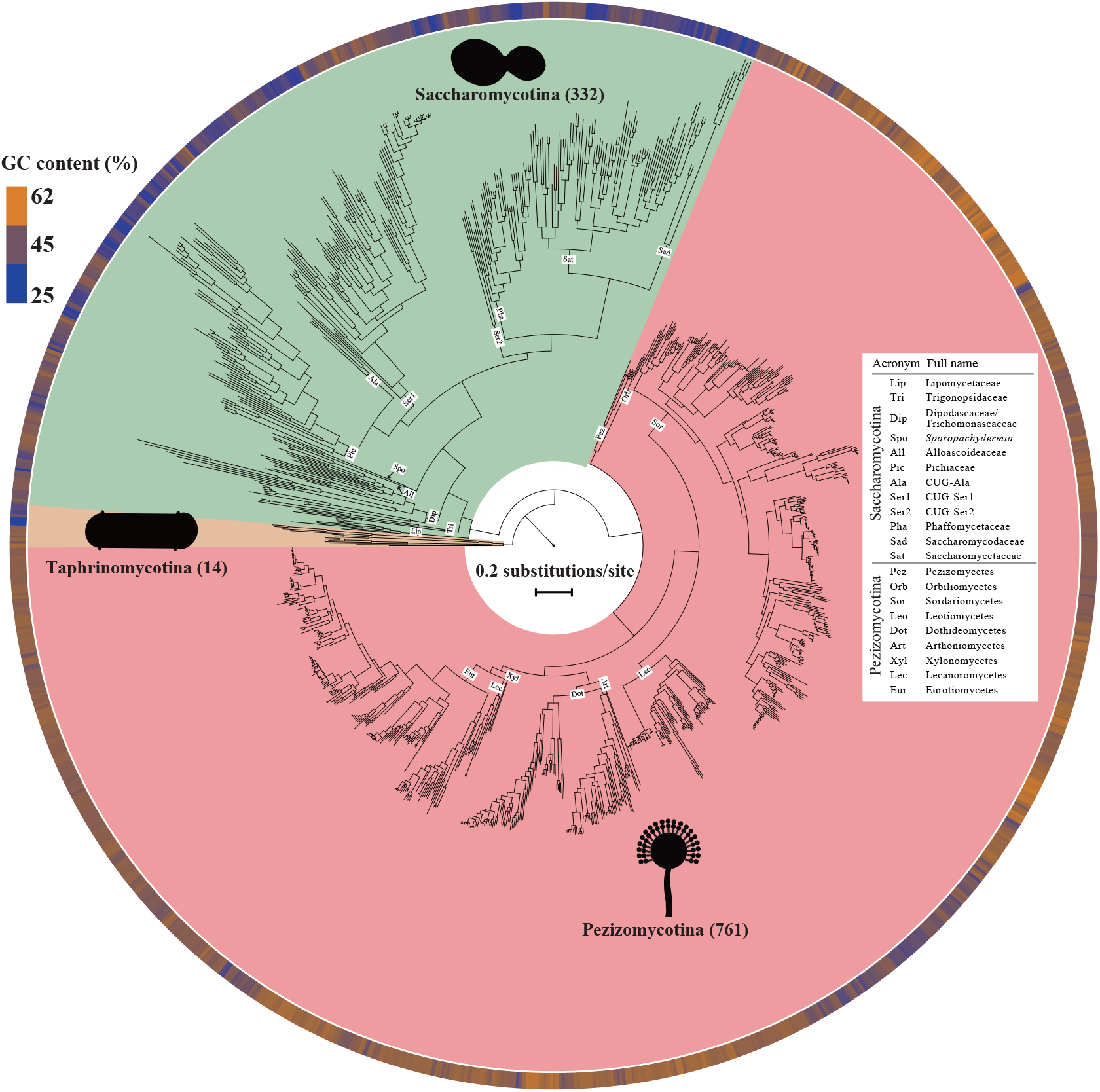
Maximum likelihood (ML) phylogeny of 1,107 taxa in the fungal phylum Ascomycota. The concatenation-based ML phylogeny (*lnL*= −269043834.145) was inferred from a set of 815 BUSCO amino acid genes (total 56, 2376 sites) under a single LG + G4 substitution model using IQ-TREE multicore version 1.5.1. The number of species sampled in each subphylum is given in parentheses. Internal branch labels are acronyms for 12 major clades in the subphylum Saccharomycotina and 9 classes in the subphylum Pezizomycotina. The bar next to each species indicates the guanine-cytosine (GC) content. On average, lineages in the subphylum Saccharomycotina have significantly lower GC content (49.6% vs. 40.6%; Wilcoxon rank-sum test; *P*-value = 3.07 x 10^-103^) but higher evolutionary rate (1.80 substitutions per site vs. 1.12 substitutions per site; Wilcoxon rank-sum test; *P*-value = 6.57 x 10^-126^) compared to lineages in the subphylum Pezizomycotina. The complete phylogenetic relationships of 1,107 taxa are given in Supplementary Fig. 2 and in the Figshare repository. For easy determination of the relationships among any subset of taxa, the phylogeny is also available through Treehouse^87^.

Our higher-level phylogeny of Ascomycota is generally more congruent with previous genome-scale phylogenies^2,18,19,35^ than with few-genes-from-many-taxa phylogenies^13–16^, particularly with respect to relationships among the nine classes in the subphylum Pezizomycotina. For example, genome-scale studies, including ours, consistently favor a clade consisting of Pezizomycetes and Orbiliomycetes as the sister group to the rest of the Pezizomycotina ^18,19^, while studies based on a few genes recovered either Orbiliomycetes^15–17^ or Pezizomycetes^14^ as the sister class to the rest of the Pezizomycotina (Fig. 2a). Our phylogeny also strongly supported the placement of the class Schizosaccharomycetes, which includes the model organism *Schizosaccharomyces pombe*, as the sister group to the class Pneumocystidomycetes, which contains the human pathogen *Pneumocystis jirovecii* (Fig. 2b). Interestingly, a recent genome-scale study of 84 fungal genomes showed that our result is consistent with the phylogeny inferred using an alignment-free composition vector approach but not with the phylogeny inferred using maximum likelihood, which instead recovered Schizosaccharomycetes as the sister group to Taphrinomycetes^18^. Finally, both concatenation- and coalescent-based approaches supported the placement of the subphylum Saccharomycotina as the sister group to the subphylum Pezizomycotina (Figs. 1 and 2c). This result is consistent with most previous studies that analyze multiple sequence alignment data^16,18,20,36–38^, but not with a recent study that analyzed genomic data with an alignment-free method and placed the subphylum Saccharomycotina as the sister group to the subphylum Taphrinomycotina^39^.

**Fig. 2.**
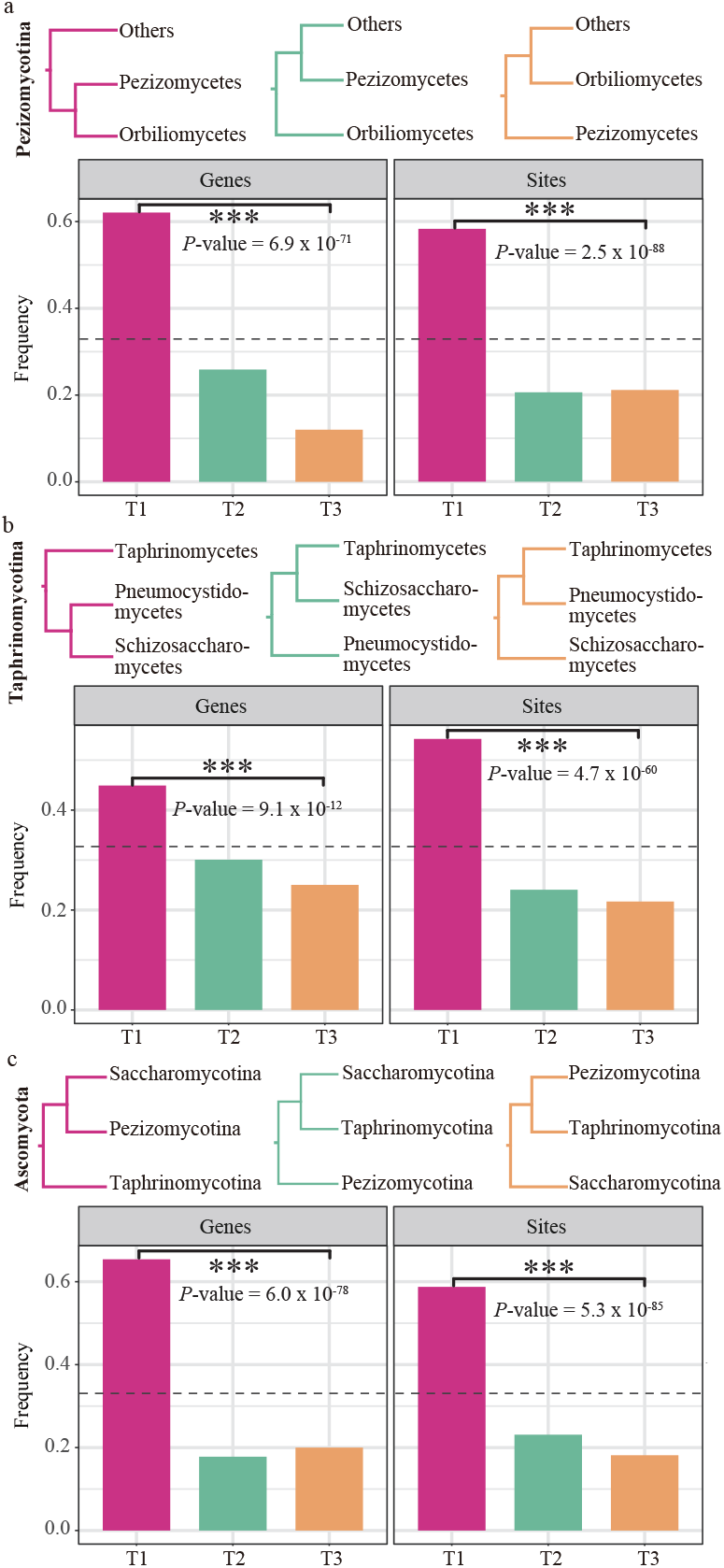
Distribution of phylogenetic signal for three historically contentious relationships within Ascomycota. For each relationship / internal branch (**a**: which class(es) is the sister group to the rest of the Pezizomycotina?; **b**: what is the relationship among three classes Schizosaccharomycetes, Pneumocystidomycetes, and Taphrinomycetes in the subphylum Taphrinomycotina?; **c**: what is the relationship among three subphyla Pezizomycotina, Saccharomycotina, and Taphrinomycotina in the phylum Ascomycota?), we applied the framework presented by Shen et al.^40^ to examine proportions of genes (left panel) and sites (right panel) supporting each of three competing hypotheses (topology 1 or T1 in red, topology 2 or T2 in green, and topology 3 or T3 in yellow). Note that both concatenation- and coalescent-based approaches supported T1 in our study. Dashed horizontal lines on 1/3 y-axis value denote expectation of proportion of genes / sites under a polytomy scenario. The *G*-test was used to test if the sets of three values are significantly different (***: *P*-value ≤ 0.001). All values are given in Supplementary Tables 4 and 5. Input and output files associated with phylogenetic signal estimation are also deposited in the Figshare repository.

To evaluate whether our genome-scale data matrix robustly resolved the three historically contentious branches discussed in the previous paragraph, we quantified the distribution of phylogenetic signal for alternative topologies of these three phylogenetic hypotheses at the level of genes and sites using a maximum likelihood framework presented by Shen et al.^40^. First, we found that phylogenetic support for each of the three branches stemmed from many genes, i.e., it was not dominated by a small number of genes with highly disproportionate influence (Supplementary Table 4). Second, we found that the topology recovered by both concatenation- and coalescent-based approaches in our study had significantly the highest frequencies of supporting genes and supporting sites (*G*–test), ranging from 0.45 to 0.65, in all three branches examined (Fig. 2a-c, Supplementary Tables 4 and 5). Importantly, none of two alternative conflicting phylogenetic hypotheses for each of the three branches received frequencies of supporting genes and supporting sites that were equal or greater than 1/3 (0.33), the value expected if the relationships among the taxa were represented by a polytomy. The very small fraction of branches where concatenation- and coalescent-based inference conflicted (<5%) and the robust support of individual genes and sites for specific historically contentious branches (Fig. 2a-c) suggest that the coupling of genome-scale amounts of data and comprehensive taxon sampling will provide robust resolution to major lineages of the tree of life^2,41^.

### A genome-scale evolutionary timetree of the fungal phylum Ascomycota

We next used the robust phylogeny, a relaxed molecular clock approach, and six widely accepted time calibration nodes (see Methods), to infer the timescale of evolution of Ascomycota. We inferred the origin of the phylum to have taken place 563 million years ago (mya) (95% credibility interval (CI): 631–495 mya); the origin of the subphylum Saccharomycotina 438.4 mya (CI: 590–304 mya); the origin of the subphylum Pezizomycotina 407.7 mya (CI: 631–405 mya); and the origin of Taphrinomycotina crown group 530.5 mya (CI: 620–417 mya). Notably, the taxonomic placement of all budding yeasts into a single class, Saccharomycetes, whose origin coincides with the origin of the subphylum Saccharomycotina, means that the last common ancestor of this sole class of budding yeasts is much more ancient than those of any of the 9 classes (based on current taxon sampling) in the subphylum Pezizomycotina (Supplementary Fig. 4 and Supplementary Table 6). For example, the most ancient class in Pezizomycotina is Pezizomycetes, whose origin is dated 247.7 mya (CI: 475-193 mya) (Supplementary Fig. 4 and Supplementary Table 6). The other outlier, albeit with much larger confidence intervals, is class Neolectomycetes in Taphrinomycotina, which we estimate to have originated 480.4 mya (CI: 607-191 mya) (Supplementary Fig. 4 and Supplementary Table 6).

Comparison of our inferred dates of divergence to those of a recent study using a 4-gene, 145-taxon data matrix^14^ shows that our estimates are younger (563 vs 588 mya for Ascomycota and 408 vs 458 mya for Pezizomycotina; sparser taxon sampling in the previous study prevents comparison of dates for Saccharomycotina and Taphrinomycotina). This result is consistent with findings of previous studies^42,43^, where inclusion of large numbers of genes was found to also result in younger estimates of divergence times, perhaps because of the influence of larger amounts of data in decreasing the stochastic error involved in date estimation. In summary, generation of a genome-scale timetree for more than 1,000 ascomycete species spanning the diversity of the phylum provides a robust temporal framework for understanding and exploring the origin and diversity of Ascomycota lifestyles^44^.

### Contrasting modes of genome evolution in fungal phylum Ascomycota

To begin understanding the similarities and differences in the modes of genome evolution between subphyla, we focused on examining the evolution of seven different genomic properties between Saccharomycotina (332 taxa) and Pezizomycotina (761 taxa), the two most taxon-rich subphyla in Ascomycota (Fig. 3). Specifically, we found that Saccharomycotina exhibited a 1.6-fold higher evolutionary rate (on average, 1.80 substitutions per site in Saccharomycotina vs. 1.12 substitutions per site in Pezizomycotina), 1.24-fold lower GC content (40% vs. 50%), 3-fold smaller genome size (13 Mb vs. 39 Mb), 1.9-fold lower number of protein-coding genes (5,734 vs. 10,847), 1.3-fold lower number of DNA repair genes (41 vs. 54), 1.2-fold higher number of tRNA genes (179 vs. 146), and 1.3-fold smaller estimates of non-synonymous to synonymous substitution rate ratio (*d*_N_/*d*_S_) (0.053 vs. 0.063), compared to Pezizomycotina (Fig. 3a, Table 1, and Supplementary Table 7).

**Fig. 3.**
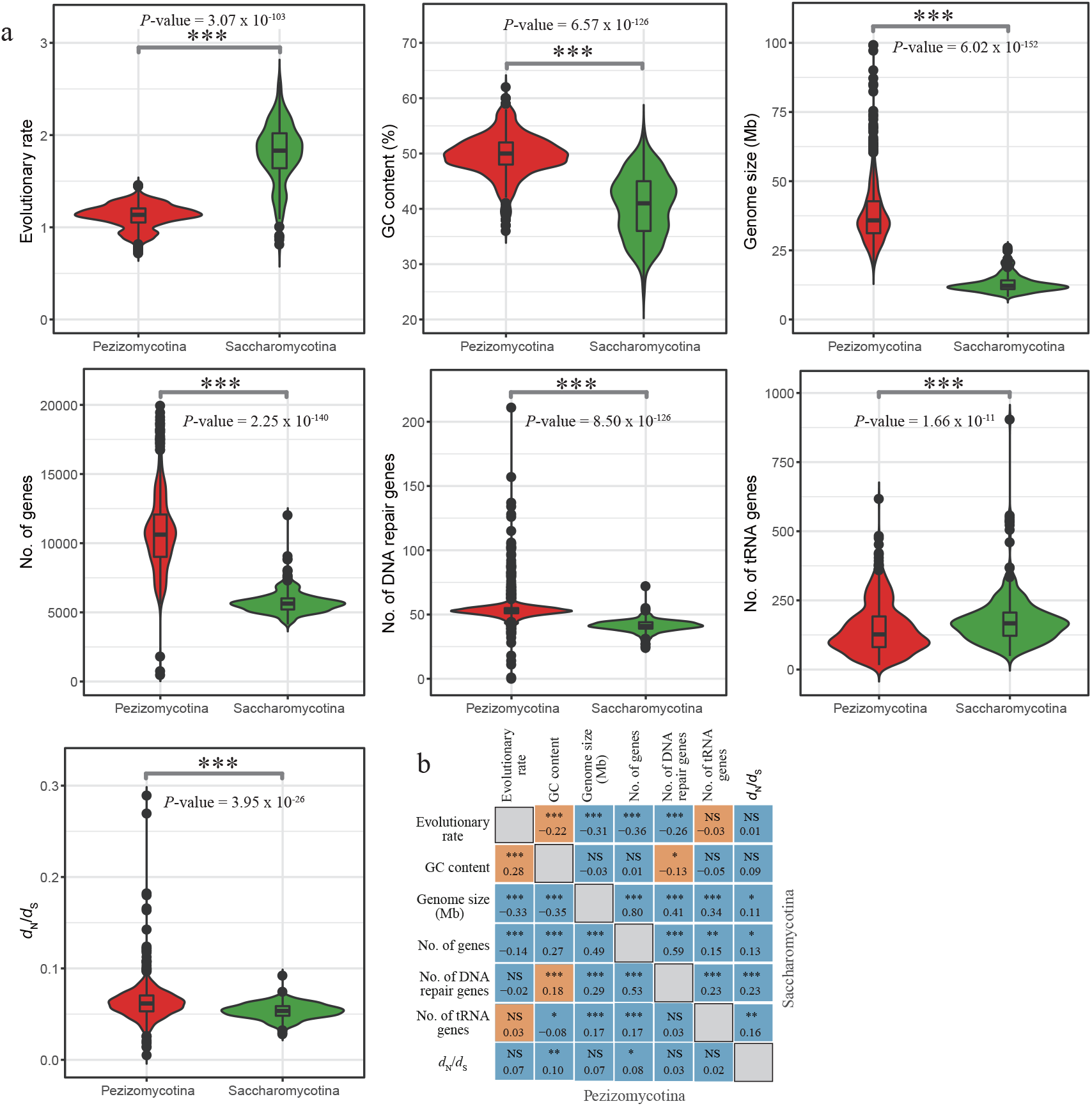
Contrasting patterns for seven genomic properties between Pezizomycotina and Saccharomycotina. **a**, For each species in Pezizomycotina (colored in red, n=761) and Saccharomycotina (colored in green, n=332), we calculated evolutionary rate, GC content, genome size, number of protein-coding genes, number of DNA repair genes, number of tRNA genes, and *d*_N_/*d*_S_ (see the Methods section for details). The Wilcoxon rank-sum test was used to test if the sets of values in two subphyla are significantly different. **b**, Pairwise standard Pearson’s correlation coefficient among pairs of the seven genomic properties were conducted using R 3.4.2 for Pezizomycotina (lower diagonal) and Saccharomycotina (upper diagonal), respectively. For each cell, the top value corresponds to *P*-value (NS: *P*-value >0.05; *: *P*-value ≤0.05; **: *P*-value ≤0.01; ***: *P*-value ≤ 0.001), whereas the bottom value corresponds to Pearson’s coefficient value. Orange cells denote instances where correlation trends in Pezizomycotina and Saccharomycotina are in opposite directions, whereas blue cells denote instances where the trends are in the same direction. The detailed values of all seven properties in Pezizomycotina and Saccharomycotina are given in Supplementary Table 7. The correlations among these seven properties are largely consistent before (i.e., standard Pearson’s correlations) and after (i.e., phylogenetically independent contrasts) accounting for correlations due to phylogeny (see Supplementary Table 8).

**Table 1.** Summary of values for seven genomic properties in extant Saccharomycotina and Pezizomycotina and in the last common ancestors of Saccharomycotina and Pezizomycotina.

| Property | Extant Saccharomycotina*<br>(n=332) | Extant Pezizomycotina*<br>(n=761) | Saccharomycotina<br>ancestor | Pezizomycotina<br>ancestor | Difference between<br>two extant lineages | Difference between<br>two ancestors |
| --- | --- | --- | --- | --- | --- | --- |
| Evolutionary rate (amino<br>acid substitutions/ site) | 1.80 | 1.12 | 1.1 | 0.9 | 0.68 | 0.2 |
| GC content (%) | 40 | 50 | 43 | 47 | 10 | 4 |
| Genome size (Mb) | 13 | 39 | 23 | 42 | 26 | 19 |
| No. of genes | 5,734 | 10,847 | 7,000 | 9,400 | 5,113 | 2,400 |
| No. of DNA repair genes | 41 | 54 | 44 | 52 | 13 | 8 |
| No. of tRNA genes | 179 | 146 | 160 | 170 | 33 | 10 |
| $d_N/d_S$ | 0.053 | 0.063 | 0.052 | 0.058 | 0.01 | 0.006 |

\* denote average values.

Analysis of standard Pearson’s correlations among the seven genomic properties revealed that two pairs exhibited statistically significant contrasting patterns between Saccharomycotina and Pezizomycotina. Specifically, evolutionary rate shows negative correlation with GC content in Saccharomycotina but positive correlation in Pezizomycotina and GC content shows negative correlation with number of DNA repair genes in Saccharomycotina but positive correlation in Pezizomycotina (Fig. 3b). These correlations are largely consistent before (i.e., standard Pearson’s correlations) and after (i.e., phylogenetically independent contrasts) accounting for correlations due to phylogeny (Supplementary Table 8).

For each of the seven properties, we used our genome-scale phylogeny (Fig. 1) to infer the ancestral character states and reconstruct their evolution in the Saccharomycotina ancestor and the Pezizomycotina ancestor. Comparison of ancestral states along branches on the Saccharomycotina part of the phylogeny to those on the Pezizomycotina part of the phylogeny shown that all genomic properties, except the number of tRNA genes, exhibited different modes of evolution (Fig. 4 and Table 1). For example, most Saccharomycotina branches exhibit evolutionary rates of at least 1.0 amino acid substitutions / site, whereas those of Pezizomycotina exhibit evolutionary rates between 0.7 and 1.4 substitutions / site (Fig. 4a). However, the inferred values for these properties in the Saccharomycotina last common ancestor and in the Pezizomycotina last common ancestor nodes are quite similar. For example, the inferred state values for the Saccharomycotina last common ancestor and the Pezizomycotina last common ancestor are 1.1 and 0.9 substitutions / site for evolutionary rate and 43% and 47% for GC content (Table 1), respectively. Interestingly, the same trends are also observed across lineages, such as Lipomycetaceae, which is the sister group to the rest of the Saccharomycotina, and the clade consisting of Pezizomycetes and Orbiliomycetes, which is the sister group to the rest of the Pezizomycotina (Fig. 4a and b).

**Fig. 4.**
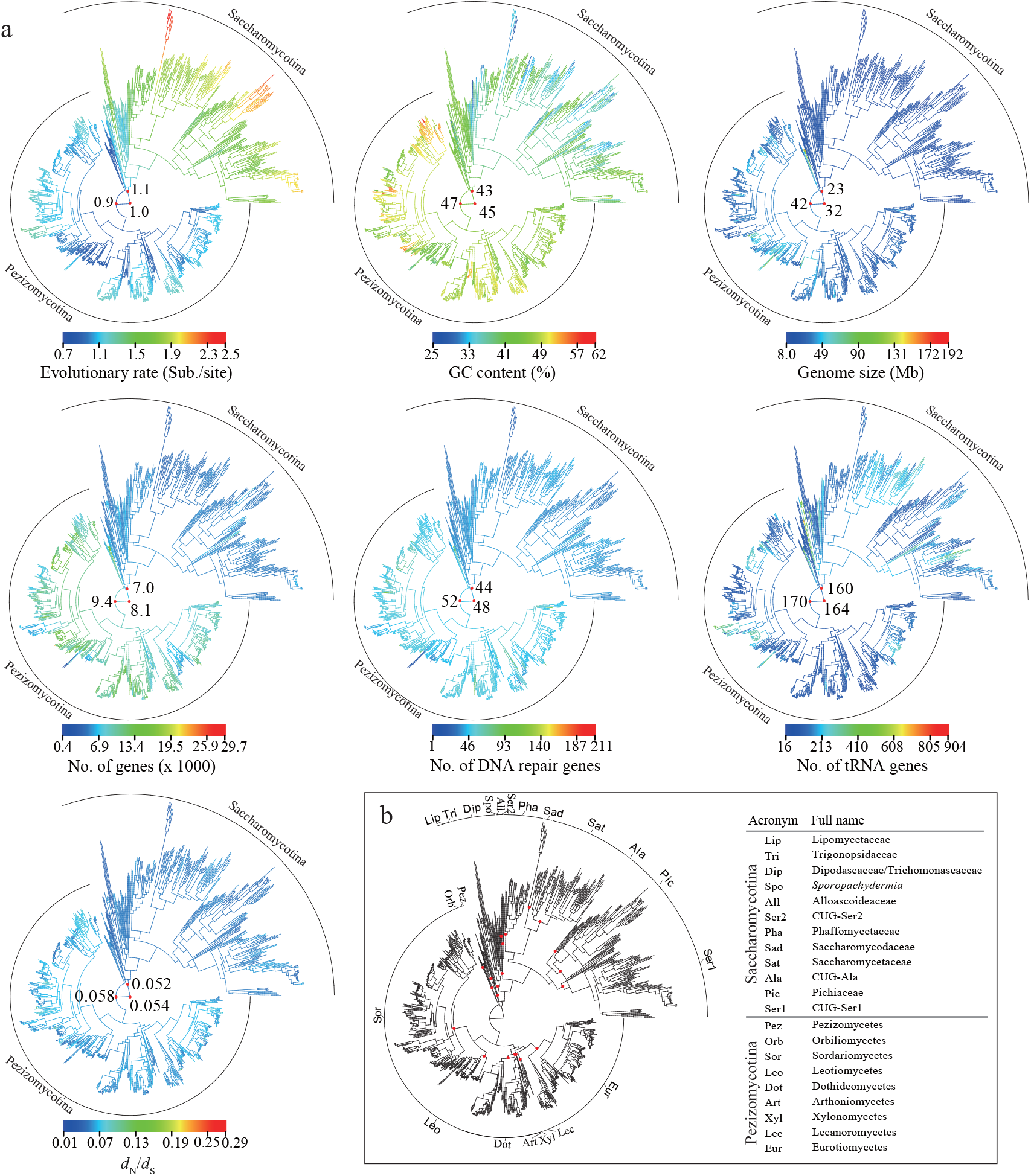
Contrasting modes of genome evolution in Pezizomycotina and Saccharomycotina. **a**, For each of the seven genomic properties examined (see the Methods section for details), we reconstructed them as continuous traits on the species phylogeny (Fig. 1) and visualized their ancestral states with the R package phytools v0.6.44 ^86^. Heatmap bars denote ancestral state values from small (blue) to large (red). Three ancestral state values next to three red dots are shown for the ancestor of the subphyla Pezizomycotina and Saccharomycotina, the ancestor of the subphylum Pezizomycotina, and the ancestor of the subphylum Saccharomycotina, respectively. **b**, Phylogeny key showing the placement of the 21 nodes representing the last common ancestors of the 12 major clades in the subphylum Saccharomycotina and of the 9 classes in the subphylum Pezizomycotina; the 21 nodes are indicated by the red dots. The orders of branches in **a** are identical to those in **b**.

Comparison of the trait values for the seven genome properties between extant Saccharomycotina and Pezizomycotina branches to those of the Saccharomycotina and Pezizomycotina last common ancestors showed that evolutionary rate, GC content, genome size, and number of protein-coding genes were the properties with the highest amounts of evolutionary change (Figs. 3 and 4, Table 1). Ancestral state reconstruction also enabled inference of the direction of evolutionary change for each of the evolutionary properties. For example, the Saccharomycotina and Pezizomycotina last common ancestors, as well as branches in Lipomycetaceae and branches across Pezizomycotina, exhibit similar evolutionary rates, whereas the rest of the nodes and branches in the Saccharomycotina part of the phylogeny exhibit much higher evolutionary rates. This pattern suggests that the higher levels of genomic diversity in Saccharomycotina stem from an acceleration of evolutionary rate that occurred within the subphylum, after the divergence of Lipomycetaceae from the rest of the Saccharomycotina (Fig. 4a and b).

Why do Saccharomycotina exhibit higher evolutionary rates compared to Pezizomycotina? Studies in other lineages, such as vertebrate^45^ and invertebrate^46^ animals, have previously shown that evolutionary rate is positively associated with generation time. Assuming that mutation rates are equal, species with shorter generation times will replicate their genomes more frequently, accruing more mutations per unit time. While the generation times of most fungi in our phylogeny are unknown, the generation times of model organisms in Saccharomycotina are thought to be shorter than those in Pezizomycotina. For example, the doubling time of the budding yeasts *S. cerevisiae* and *C. albicans* under optimal conditions is 90 min^47,48^, while that of the filamentous fungi *Aspergillus nidulans* and *Neurospora crassa* is between 2-3 hours^49,50^. An alternative but not mutually exclusive explanation may be that Saccharomycotina have, on average, 13 fewer DNA repair genes (41) than Pezizomycotina (54) (Fig. 3 and Table 1), since it is well established that absence or loss of DNA repair genes increase mutation rates^51–53^. The lower numbers of DNA repair genes in budding yeasts, but not their higher evolutionary rate, was also recently reported in a recent analysis of 328 ascomycete proteomes by Milo et al.^54^. Finally, other life-history traits (e.g., smaller cell size, faster metabolism, and larger population size) that have been associated with variation in the rate of molecular evolution^55^ might also contribute to higher evolutionary rates of the Saccharomycotina.

Variation in genomic GC content has historically been of broad interest in biology^56^. Average GC content values of different genomic regions (e.g., intergenic regions, protein-coding regions) in Saccharomycotina are consistently lower than those in Pezizomycotina (Supplementary Fig. 5). Similarly, gene-wise average estimates of GC content showed that all 815 BUSCO genes in Saccharomycotina have lower GC content values than those in Pezizomycotina (Supplementary Fig. 6). Moreover, we found that the frequencies of amino acids encoded by GC-rich codons in Saccharomycotina are much lower than those of amino acids encoded by GC-rich codons in Pezizomycotina (Supplementary Fig. 7). Ancestral stat ereconstruction of genomic GC content along branches on the phylogeny shows that the Saccharomycotina and Pezizomycotina last common ancestors, as well as branches in Lipomycetaceae and branches in classes Pezizomycetes and Orbiliomycetes, exhibit intermediate GC content around 45%. In contrast, GC content of most branches within the rest of Saccharomycotina (i.e., all major clades of Saccharomycotina, including extant taxa, except Lipomycetaceae) evolved toward 40%, while GC content within the rest of Pezizomycotina (i.e., all classes, including extant taxa, except Pezizomycetes and Orbiliomycetes) evolved toward 50%. This pattern suggests that the evolution of lower levels of GC content in Saccharomycotina occurred after the divergence of Lipomycetaceae from the rest of Saccharomycotina and that the evolution of higher levels of GC content in Pezizomycotina occurred after the divergence of the clade consisting of Pezizomycetes and Orbiliomycetes from the rest of Pezizomycotina (Fig. 4a and b).

Why are Pezizomycotina genomes more GC-rich compared to Saccharomycotina genomes? There are two possible explanations. The first one is that mutational biases have skewed the composition of Saccharomycotina genomes toward AT content^57^. For example, Steenwyk et al. showed that *Hanseniaspora* budding yeasts with higher AT content lost a greater number of DNA repair genes than those with lower AT content^53^, suggesting that the loss of DNA repair genes is associated with AT richness. Consistent with these results, we found that Pezizomycotina genomes contain a higher number of DNA repair genes than Saccharomycotina (Fig. 3 and Table 1). The second potential, not necessarily mutually exclusive, explanation is that mutational biases have skewed Pezizomycotina genomes toward GC richness. It was recently shown that increasing GC-biased gene conversion (gBGC), a process associated with recombination that favors the transmission of GC alleles over AT alleles^58^, can result in a systematic underestimate of *d*_N_/*d*_S_ in birds^59^. If this is true for Ascomycota, due to the higher GC content of Pezizomycotina genomes, we would expect that their *d*_N_/*d*_S_ would be underestimated due to the higher levels of gBGC compared to Saccharomycotina. Consistent with this expectation, by calculating differences in *d*_N_/*d*_S_ before and after accounting for gBGC across 815 codon-based BUSCO genes, we found that the underestimate of *d*_N_/*d*_S_ in Pezizomycotina is 2-fold higher than that in Saccharomycotina (Pezizomycotina: average of differences in *d*_N_/*d*_S_ = 0.004; Saccharomycotina: average of differences in *d*_N_/*d*_S_ = 0.002) (Supplementary Fig. 8).

## Concluding Remarks

In this study, we took advantage of the recent availability of the genome sequences of 1,107 Ascomycota species from Saccharomycotina (332), Pezizomycotina (761), and Taphrinomycotina (14) to infer a genome-scale phylogeny and timetree for the entire phylum and compare the mode of genome evolution across its subphyla. Leveraging genome-scale amounts of data from the most comprehensive taxon set to date enabled us to test the robustness of our inference for several contentious branches, potentially resolving controversies surrounding key higher-level relationships within the Ascomycota phylum. For example, our study robustly supported Saccharomycotina as the sister group to

Pezizomycotina and a clade comprised of classes Pezizomycetes and Orbiliomycetes as the sister group to the rest of the Pezizomycotina. Our first genome-scale timetree suggests the last common ancestor of Ascomycota likely originated in the Ediacaran period. Examination of mode of genome evolution revealed that Saccharomycotina, which contains the single currently described class Saccharomycetes, and Pezizomycotina, which contains 13 classes, exhibited greatly contrasting evolutionary processes for seven genomic properties, in particular for evolutionary rate, GC content, and genome size. Our results provide a robust evolutionary framework for understanding the diversification of the largest fungal phylum.

## Methods

### Data collection

To collect the greatest possible set of genome representatives of the phylum Ascomycota as of 14 December, 2018, we first retrieved the 332 publicly available Saccharomycotina yeast genomes (https://doi.org/10.6084/m9.figshare.5854692) from a recent comprehensive genomic study of the Saccharomycotina yeasts^2^. We then used “Pezizomycotina” and “Taphrinomycotina” as search terms in NCBI’s Genome Browser (https://www.ncbi.nlm.nih.gov/genome/browse#!/eukaryotes/Ascomycota) to obtain the basic information of strain name, assembly accession number, assembly release date, assembly level (e.g., contig, scaffold, etc.), and GenBank FTP access number for draft genomes from the subphyla Pezizomycotina and Taphrinomycotina, respectively. For species with multiple isolates sequenced, we only included the genome of the isolate with the highest assembly level and the latest release date. We next downloaded genome assemblies from GenBank data via FTP access number (ftp://ftp.ncbi.nlm.nih.gov/genomes/). Collectively, we included 332 species representing all 12 major clades of the subphylum Saccharomycotina^2^, 761 species representing 9 / 13 classes of the subphylum Pezizomycotina^1,6^, and 14 species representing 4 / 5 classes of the subphylum Taphrinomycotina^1,6^. Finally, we used the genomes of six representatives of the phylum Basidiomycota as outgroups. Detailed information of taxonomy and source of the 1,113 genomes in our study is provided in Supplementary Tables 1 and 2.

### Assessment of genome assemblies and phylogenomic data matrix construction

To assess the quality of each of the 1,113 genome assemblies, we used the Benchmarking Universal Single-Copy Orthologs (BUSCO), version 3.0.2^34^. Each assembly’s completeness was assessed based on the presence / absence of a set of 1,315 predefined orthologs (referred to as BUSCO genes) from 75 genomes in the OrthoDB Version 9 database^60^ from the Ascomycota database, as described previously^28,61^. In brief, for each BUSCO gene, its consensus orthologous protein sequence among the 75 reference genomes was used as query in a tBLASTn search against each genome to identify up to three putative genomic regions, and the gene structure of each putative genomic region was predicted by AUGUSTUS v 3.2.2^62^. Next, the sequences of these predicted genes were aligned to the HMM-profile of the BUSCO gene. BUSCO genes in a given genome assembly were considered as single-copy, “full-length” if there was only one complete predicted gene present in the genome, duplicated, “full-length” if there were two or more complete predicted genes present in the genome, “fragmented” if the predicted gene was shorter than 95% of the aligned sequence lengths from the 75 reference species, and “missing” if there was no predicted gene present in the genome.

To construct the phylogenomic data matrix, we started with the set of 1,315 single-copy, fulllength BUSCO genes from 1,107 representatives of the phylum Ascomycota and six outgroups. For each BUSCO gene, we first translated nucleotide sequences into amino acid sequences, taking into account the different usage of the CUG codon in Saccharomycotina^2,63^. Next, we aligned the amino acid sequences using MAFFT v7.299b^64^ with the options “--thread 4 --auto --maxiterate 1000” and trimmed amino acid alignments using the trimAl v1.4.rev15^65^ with the options “-gappyout -colnumbering”. We mapped the nucleotide sequences on the trimmed amino acid alignment based on the column numbers in the original alignment and to generate the trimmed codon-based nucleotide alignment. Finally, we removed BUSCO gene alignments whose taxon occupancy (i.e., percentage of taxa whose sequences were present in the trimmed amino acid alignment) was < 50% for each subphylum (i.e., < 7 Taphrinomycotina, < 166 Saccharomycotina, and < 381 Pezizomycotina) or whose trimmed alignment length was < 300 amino acid sites. These filters resulted in the retention of 815 BUSCO gene alignments, each of which had ≥ 50% taxon occupancy for each subphylum and alignment length ≥ 300 amino acid sites.

### Phylogenetic analysis

For each of 815 BUSCO genes, we first inferred its best-fitting amino acid substitution model using IQ-TREE multi-thread version 1.6.8 ^66^ with options “-m TEST -mrate G4” with the Bayesian information criterion (BIC). We then inferred best-scoring maximum likelihood (ML) gene tree under 10 independent tree searches using IQ-TREE. The detailed parameters for running each gene were kept in log files (see the Figshare repository). We inferred the concatenation-based ML tree using IQ-TREE on a single node with 32 logical cores under a single “LG +G4” model with the options “-seed 668688 -nt 32 -mem 220G -m LG+G4 -bb 1000”, as 404 out of 815 genes favored “LG +G4”^67,68^ as best-fitting model (see Supplementary Table 3). We also inferred the coalescent-based species phylogeny with ASTRAL-III version 4.10.2^69,70^ using the set of 815 individual ML gene trees. The reliability of each internal branch was evaluated using 1,000 ultrafast bootstrap replicates^71^ and local posterior probability^72^, in the concatenation- and coalescence-based species trees, respectively. We visualized phylogenetic trees using the R package *ggtree* v1.10.5^73^.

We used the non-Bayesian RelTime method, as implemented in the command line version of MEGA7^74^ to estimate divergence times. The very large size of our data matrix, both in terms of genes as well as in terms of taxa, prohibited the use of computationally much more demanding methods, such as the Bayesian MCMCTree method^75,76^. The concatenation-based ML tree with branch lengths was used as the input tree. Six time calibration nodes, which were retrieved from the TimeTree database ^77^, were used for molecular dating analyses: the *Saccharomyces cerevisiae* – *Saccharomyces uvarum* split (14.3 mya – 17.94 mya), the *Saccharomyces cerevisiae* – *Kluyveromyces lactis* split (103 mya – 126 mya), the *Saccharomyces cerevisiae* – *Candida albicans* split (161 mya – 447 mya), the origin of the subphylum Saccharomycotina (304 mya – 590 mya), the *Saccharomyces cerevisiae* – *Saitoella complicata* split (444 mya – 631 mya), and the origin of the subphylum Pezizomycotina (at least 400 mya) based on the *Paleopyrenomycites devonicus* fossil^78^.

### Examination of seven genome properties

As the subphylum Taphrinomycotina (No. species = 14) has a much smaller number of species than the subphylum Saccharomycotina (No. species = 332) and the subphylum Pezizomycotina (No. species = 761) in our dataset, we focused our analyses on the comparisons of seven genome properties (evolutionary rate, GC content, genome size, number of genes, number of DNA repair genes, number of tRNA genes, and *d*_N_/ *d*_S_) between Saccharomycotina and Pezizomycotina. Specifically, for a given taxon, 1) evolutionary rate is a sum of path distances from the most common ancestor of the subphyla Saccharomycotina and Pezizomycotina to its tip on the concatenation-based ML tree (Fig. 1); 2) GC content is the percentage of guanine-cytosine nucleotides in genome; 3) genome size is the total number of base pairs in genome in megabases (Mb); 4) number of genes is the number of proteincoding genes in genome. The gene structure was predicted with AUGUSTUS v3.3.1 ^79^ on *Aspergillus fumigatus* and *Saccharomyces cerevisiae* S288C trained models for Pezizomycotina and Saccharomycotina, respectively; 5) number of DNA repair genes was estimated by counting the number of unique protein-coding genes with GO terms related to DNA repair using InterProscan version 5 ^80^; 6) number of tRNA genes is the number of tRNA genes inferred to be present using the tRNAscan-SE 2.0 program^81^; and 7) *d*^N^*d*^S^ was estimated by calculating the average of the ratio of the expected numbers of non-synonymous *(dri)* and synonymous substitutions *(dS)* across 815 trimmed codon-based BUSCO gene alignments under the YN98 (F3X4)^82^ codon model and the free ratio model using bppml and MapNH in the bio++ libraries^83^, following the study by Bolívar et al.^59^.

### Statistical analyses

All statistical analyses were performed in R v. 3.4.2 (R core team 2017). Pearson’s correlation coefficient was used to test for correlations among seven variables. To account for phylogenetic relationships of species in correlation analysis, we used the R package ape v5.1^84^ in order to compute phylogenetically independent contrasts following the method described by Felsenstein^85^.

### Ancestral state reconstruction

To reconstruct ancestral character states for each of seven continuous properties, we used the R package phytools v0.6.44 function *contMap* ^86^ to infer ancestral character states across internal nodes using the maximum likelihood method with the function *fastAnc* and to interpolate the states along each edge using equation [2] of Felsenstein^85^. The input tree was derived from the concatenation-based ML with branch lengths, which was then pruned to keep the 1,093 taxa from the subphyla Pezizomycotina and Saccharomycotina.

## Data availability

All genome assemblies and proteomes are publicly available in the Zenodo repository: https://doi.org/10.5281/zenodo.3783970. Multiple sequence alignments, phylogenetic trees, trait ancestral character state reconstructions, log files, R codes, and custom Perl scripts are available on the figshare repository (https://doi.org/10.6084/m9.figshare.12196149; https://figshare.com/articles/Phylogenomics_and_contrasting_modes_of_genome_evolution_in_Ascomycota/12196149 – please note that this link will become active upon publication).

## Acknowledgments

We thank members of the Rokas and Hittinger labs, especially members of the Y1000+ Project for constructive feedback.

## Funding Statement

This work was conducted in part using the resources of the Advanced Computing Center for Research and Education (ACCRE) at Vanderbilt University. X.X.S. was supported by the start-up grant from the “Hundred Talents Program” at Zhejiang University and the Fundamental Research Funds for the Central Universities (No. is appending). X.Z. was supported by the National Key Project for Basic Research of China (973 Program, No. 2015CB150600) and the open fund from Key Laboratory of Ministry of Education for Genetics, Breeding and Multiple Utilization of Crops, College of Crop Science, Fujian Agriculture and Forestry University (GBMUC-2018-005). M.G. was supported by the Royal Netherlands Academy of Arts and Sciences. C.T.H. was supported by the National Science Foundation (DEB-1442148), the USDA National Institute of Food and Agriculture (Hatch Project No. 1020204), in part by the DOE Great Lakes Bioenergy Research Center (DOE BER Office of Science No. DE-SC0018409), the Pew Charitable Trusts (Pew Scholar in the Biomedical Sciences), and the Office of the Vice Chancellor for Research and Graduate Education with funding from the Wisconsin Alumni Research Foundation (H. I. Romnes Faculty Fellow). J.L.S. and A.R. were supported by the Howard Hughes Medical Institute through the James H. Gilliam Fellowships for Advanced Study program. A.R. was supported by the National Science Foundation (DEB-1442113), the Guggenheim Foundation, and the Burroughs Wellcome Fund.

## Author contributions

Study conception and design: X.X.S., C.T.H., A.R..; Acquisition of data: X.X.S.; Analysis and interpretation of data: X.X.S., J.L.S., A.L.L., D.A.O., X.Z., J.K., Y.L., M.G., C.T.H., A.R..; Drafting of manuscript: X.X.S., A.R.; Critical revision: all authors.

## Competing interests

The authors declare no competing financial interests.

## Notes

### Competing Interest Statement

The authors have declared no competing interest.

