## Supplementary Figures for "Genome-scale phylogeny and contrasting modes of genome evolution in the fungal phylum Ascomycota"

### Supplementary information

This file contains Supplementary Figs. 1-8.

**Supplementary Table 1 The taxonomic distribution of 1,107 genomes from the phylum Ascomycota based on the NCBI taxonomy.**

**Supplementary Table 2 Taxonomy and source information of 1,107 taxa in the phylum Ascomycota and six outgroups in the phylum Basidiomycota.**

**Supplementary Table 3 Summary information of the 815 BUSCO genes used in our evolutionary analyses.**

**Supplementary Table 4 Gene-wise phylogenetic signal for three contentious relationships in the Ascomycota phylum.**

**Supplementary Table 5 Site-wise phylogenetic signal for three contentious relationships in the Ascomycota phylum.**

**Supplementary Table 6 Detailed results of divergence time estimates among 4 classes in Taphrinomycotina, 9 classes in Pezizomycotina, and 12 major clades in Saccharomycotina.**

**Supplementary Table 7 All values of genomic completeness and seven properties for each of 1,113 taxa.**

**Supplementary Table 8 All correlations between seven properties before (Standard correlation coefficient) and after (Phylogenetically independent contrasts) taking phylogenetic relationships of species into account in Saccharomycotina and Pezizomycotina, respectively.**

**Supplementary Figure 1. The distribution of genomic completeness of 1,107 genomes from the phylum Ascomycota.** Genomic completeness was assessed as the percentage of the full-length genes based on a set of the 1,315 Benchmarking Universal Single-Copy Orthologs (BUSCO). Exact values of the 1,107 genomes are provided in Supplementary Table 7.

**Supplementary Figure 2. The phylogenetic relationships of 1,107 taxa in the Ascomycota and six outgroups inferred from the concatenation-based analysis of 815 BUSCO amino acid genes (total 56, 2376 sites).** The concatenation-based ML phylogeny (*lnL*= -269043834.145) was reconstructed under a single LG + G4 substitution model using IQ-TREE multicore version 1.5.1. Branch support values near internodes / internal branches correspond to ultrafast bootstrap support. Only support values smaller than 95% are shown. The 46 branches shown in red denote conflicts between the concatenation-based phylogeny and the coalescence-based phylogeny (Supplementary Fig. 3). The higher-level phylogenetic relationships among major lineages in this phylogeny are shown at the upper right inset. Species names in the figure are searchable. This phylogeny is also provided in Newick format in the Figshare repository and available through Treehouse.

**Supplementary Figure 3. The phylogenetic relationships of 1,107 taxa in the Ascomycota and six outgroups inferred from coalescence-based analysis.** The coalescence-based phylogeny was reconstructed based on a set of 815 best-scoring genes trees using ASTRAL-II version 4.10.2. Branch support values near internodes correspond to local posterior probability. Only support values smaller than 95% are shown. The 46 internal branches in red denote conflicts between the concatenation-based phylogeny (Supplementary Fig. 2) and the coalescence-based phylogeny. The higher-level phylogenetic relationships among major lineages in this phylogeny are shown at the bottom left inset. Species names in the figure are searchable. This phylogeny is also provided in Newick format in the Figshare repository.

**Supplementary Figure 4. The Time-calibrated phylogeny of 1,107 taxa in the Ascomycota. a**, Higher-level timetree of 25 major groups. Number of species sampled in each group is given in parentheses; the size of the triangle is proportional to number of species. mya, million years ago; Edi., Ediacaran; Cam., Cambrian; Ord., Ordovician; Sil., Silurian; Dev., Devonian; Car., Carboniferous; Per., Permian; Tri., Triassic; Jur., Jurassic; Cre., Cretaceous; Pal., Paleogene; Neo., Neogene. The numbers near internodes / internal branches are divergence time estimates; 1 time unit = 100 million years ago (mya). **b**, the complete timetree of 1,107 taxa. Species names in the figure are searchable. This timetree is provided in Newick format in the Figshare repository.

**Supplementary Figure 5. Average GC content (%) values for different genomic regions in Saccharomycotina and Pezizomycotina.** “All”: the entire genome; “IGR”: the intergenic region; “CDS”: the coding sequence; “Pos1”: all first codon positions from all coding sequences in the genome; “Pos2”: all second codon positions; “Pos3”: all third codon positions. The gene structure of each putative genomic region was predicted with AUGUSTUS v3.3.1 using *Saccharomyces cerevisiae* S288C and *Aspergillus fumigatus* trained models for Saccharomycotina and Pezizomycotina, respectively. Overall, Saccharomycotina have lower GC content values than Pezizomycotina for every type of genomic region.

**Supplementary Figure 6. Scatterplot of gene-wise GC content values between Pezizomycotina and Saccharomycotina.** Each point corresponds to one BUSCO gene (the total number of BUSCO genes used was 815). For each BUSCO gene, the average of GC content values across Pezizomycotina (or Saccharomycotina) on codon-based nucleotide alignments was considered as the gene-wise GC content. Dashed line denotes x=y. Our results show that all 815 BUSCO genes in Saccharomycotina have lower GC content values than their orthologs in Pezizomycotina.

**Supplementary Figure 7. Difference in average amino acid frequency between Pezizomycotina and Saccharomycotina**. For each amino acid, its difference in the average amino acid frequency between Pezizomycotina and Saccharomycotina was calculated on the complete proteomes, which were predicted with AUGUSTUS v3.3.1 using *Aspergillus fumigatus* and *Saccharomyces cerevisiae* S288C trained models for Pezizomycotina and Saccharomycotina, respectively. Red bars denote amino acids whose average frequency is higher in Pezizomycotina than in Saccharomycotina, whereas green bars denote amino acids whose average frequency is higher in Saccharomycotina than in Pezizomycotina. This plot shows that Pezizomycotina genomes generally contain higher frequencies of amino acids encoded by GC-rich codons but lower frequencies of amino acids encoded by AT-rich codons compared to Saccharomycotina.

**Supplementary Figure 8. The impact of GC-biased gene conversion (gBGC) on $d_N/d_S$ estimates.** To investigate the effect of GC content on selection acting on protein-coding genes, we used an approach that accounts for non-stationary base composition, to calculate difference in $d_N/d_S$ before and after accounting for gBGC for 815 codon-based BUSCO genes in Pezizomycotina and Saccharomycotina, respectively. This plot shows that Pezizomycotina with higher GC content would suffer from greater effect of gBGC on estimating $d_N/d_S$ than Saccharomycotina with lower GC content. Calculating differences in $d_N/d_S$ after and before accounting for gBGC across 815 codon-based BUSCO genes revealed that the effect of gBGC on $d_N/d_S$ estimates in Pezizomycotina is 2-fold higher than that in Saccharomycotina

(Pezizomycotina: average of differences in $d_N/d_S$ = 0.004; Saccharomycotina: average of differences in $d_N/d_S$ = 0.002).

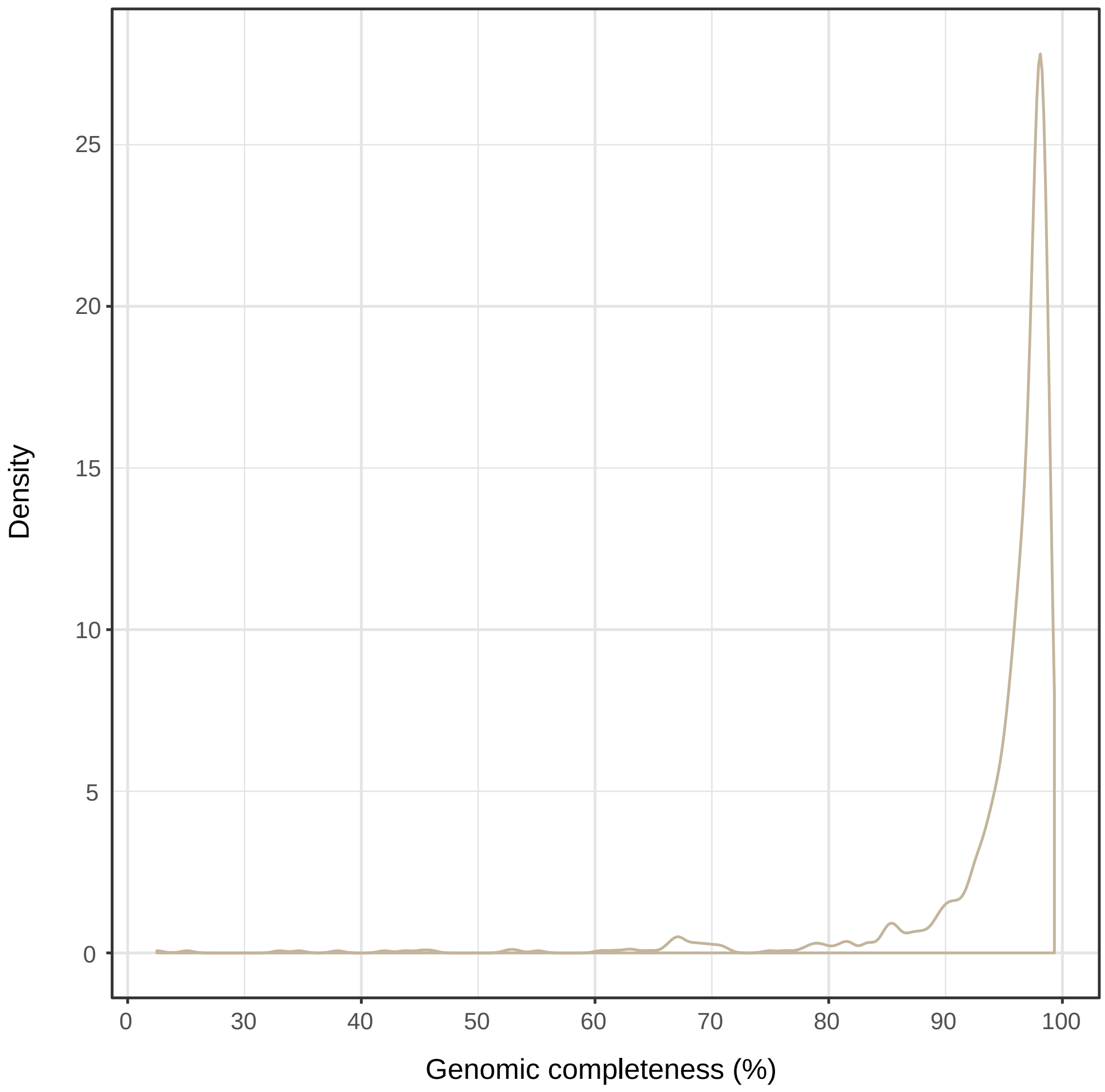


**Supplementary Figure 1. The distribution of genomic completeness of 1,107 genomes from the phylum Ascomycota.** Genomic completeness was assessed as the percentage of the full-length genes based on a set of the 1,315 Benchmarking Universal Single-Copy Orthologs (BUSCO). Exact values of the 1,107 genomes are provided in Supplementary Table 7.

**Supplementary Figure 2. The phylogenetic relationships of 1,107 taxa in the Ascomycota and 6 outgroups inferred from the concatenation-based analysis of 815 BUSCO amino acid genes (total 56, 2376 sites).** The concatenation-based ML phylogeny (*lnL*= -269043834.145) was reconstructed under a single LG + G4 substitution model using IQ-TREE multicore version 1.5.1. Branch support values near internodes / internal branches correspond to ultrafast bootstrap support. Only support values smaller than 95% are shown. The 46 branches shown in red denote conflicts between the concatenation-based phylogeny and the coalescence-based phylogeny (Supplementary Fig. 3). The higher-level phylogenetic relationships among major lineages in this phylogeny are shown at the upper right inset. Species names in the figure are searchable. This phylogeny is also provided in Newick format in the Figshare repository and available through Treehouse.

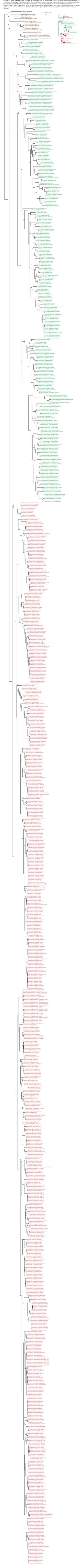

**Supplementary Figure 3. The phylogenetic relationships of 1,107 taxa in the Ascomycota and 6 outgroups inferred from coalescence-based analysis.** The coalescence-based phylogeny was reconstructed based on a set of 815 best-scoring genes trees using ASTRAL-II version 4.10.2. Branch support values near internodes correspond to local posterior probability. Only support values smaller than 95% are shown. The 46 internal branches in red denote conflicts between the concatenation-based phylogeny (Supplementary Fig. 2) and the coalescence-based phylogeny. The higher-level phylogenetic relationships among major lineages in this phylogeny are shown at the bottom left inset. Species names in the figure are searchable. This phylogeny is also provided in Newick format in the Figshare repository.

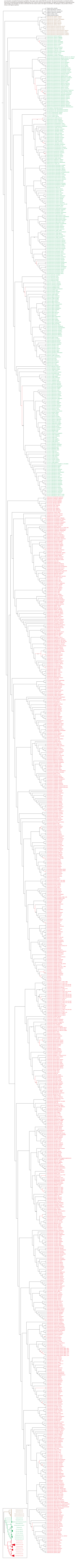

**Supplementary Figure 4. The Time-calibrated phylogeny of 1,107 taxa in the Ascomycota.** **a**, Higher-level timetree of 25 major groups. Number of species sampled in each group is given in parentheses; the size of the triangle is proportional to number of species. mya, million years ago; Edi., Ediacaran; Cam., Cambrian; Ord., Ordovician; Sil., Silurian; Dev., Devonian; Car., Carboniferous; Per., Permian; Tri., Triassic; Jur., Jurassic; Cre., Cretaceous; Pal., Paleogene; Neo., Neogene. The numbers near internodes / internal branches are divergence time estimates; 1 time unit = 100 million years ago (mya). **b**, the complete timetree of 1,107 taxa. Species names in the figure are searchable. This timetree is provided in Newick format in the Figshare repository.

**a**

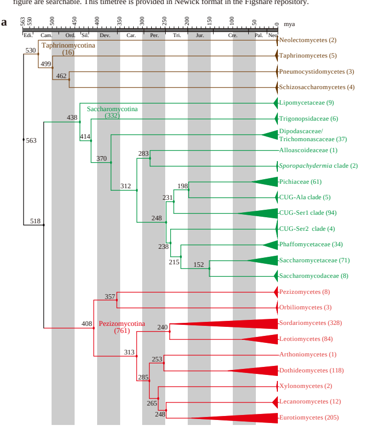


**b**

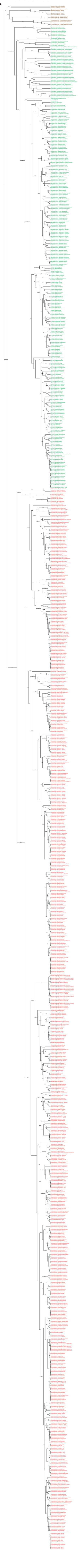

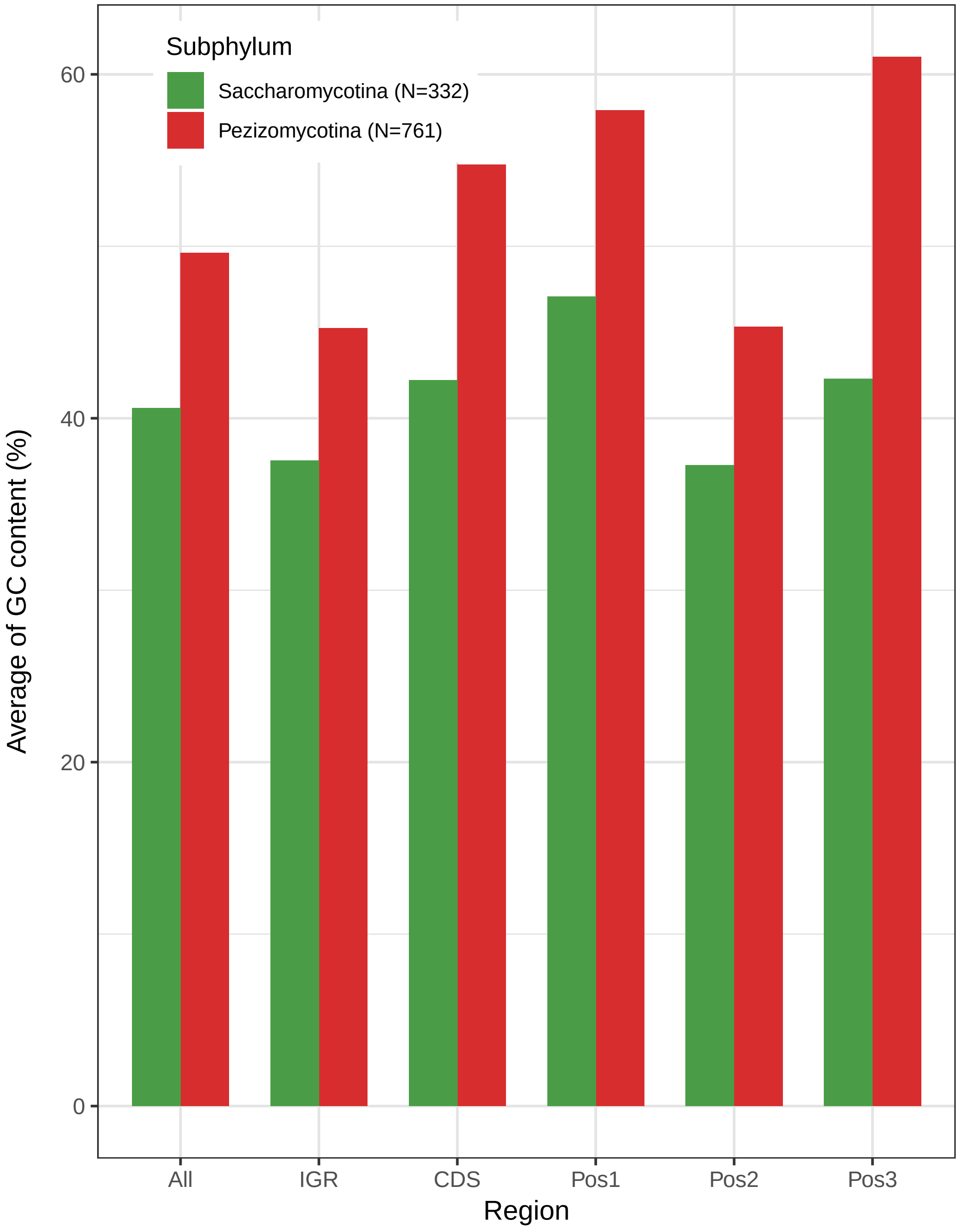


**Supplementary Figure 5. Average GC content (%) values for different genomic regions in Saccharomycotina and Pezizomycotina.** "All": the entire genome; "IGR": the intergenic region; "CDS": the coding sequence; "Pos1": all first codon positions from all coding sequences in the genome; "Pos2": all second codon positions; "Pos3": all third codon positions. The gene structure of each putative genomic region was predicted with AUGUSTUS v3.3.1 using *Saccharomyces cerevisiae* S288C and *Aspergillus fumigatus* trained models for Saccharomycotina and Pezizomycotina, respectively. Overall, Saccharomycotina have lower GC content values than Pezizomycotina for every type of genomic region.

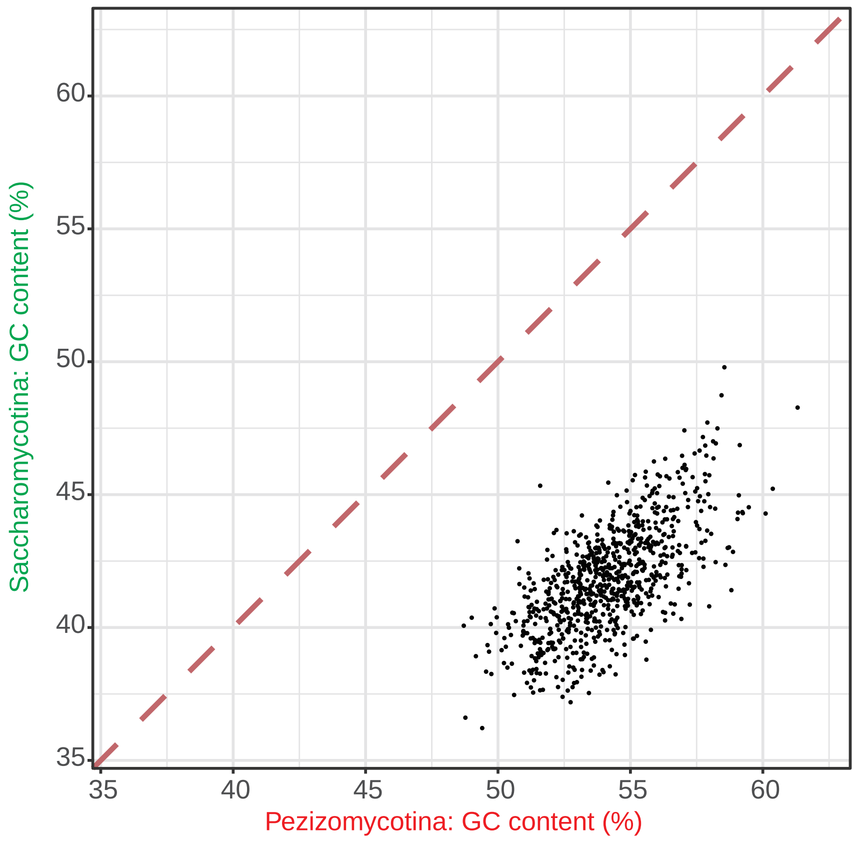


**Supplementary Figure 6. Scatterplot of gene-wise GC content values between Pezizomycotina and Saccharomycotina.** Each point corresponds to one BUSCO gene (the total number of BUSCO genes used was 815). For each BUSCO gene, the average of GC content values across Pezizomycotina (or Saccharomycotina) on codon-based nucleotide alignments was considered as the gene-wise GC content. Dashed line denotes x=y. Our results show that all 815 BUSCO genes in Saccharomycotina have lower GC content values than their orthologs in Pezizomycotina.

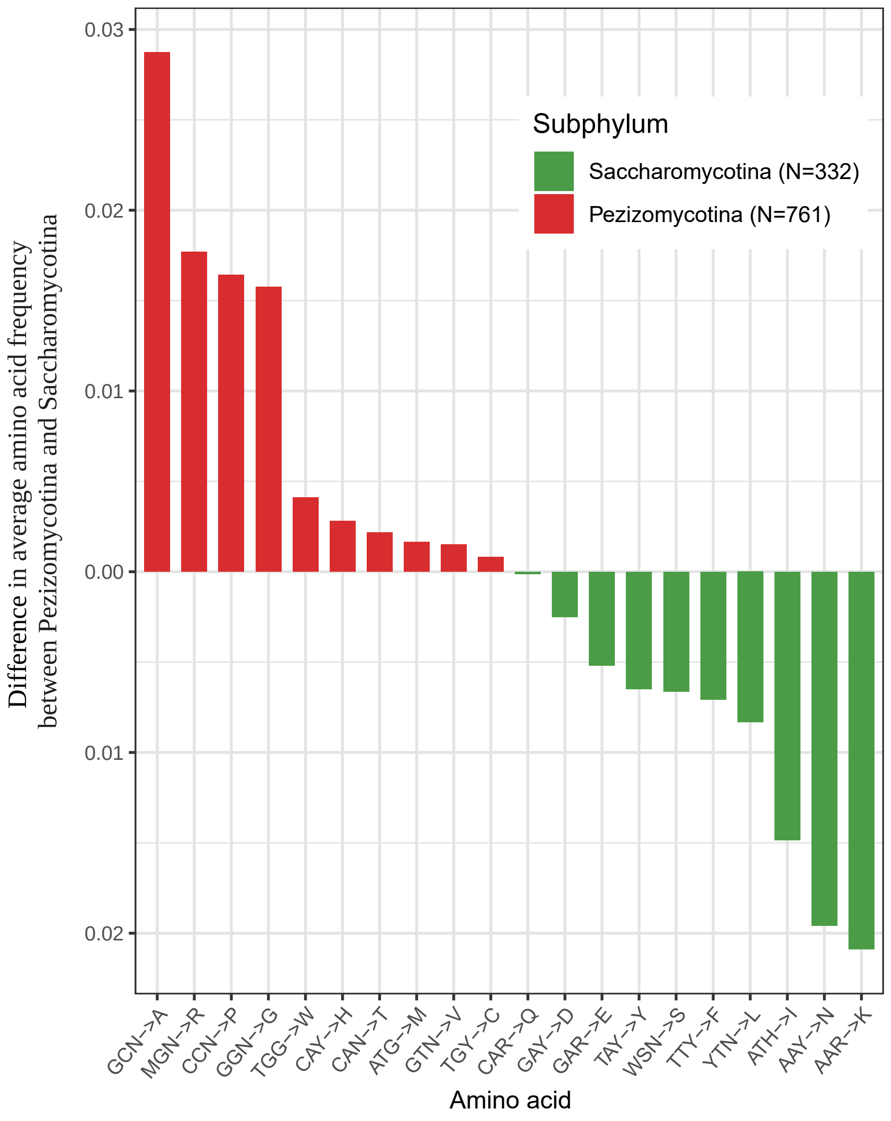


**Supplementary Figure 7. Difference in average amino acid frequency between Pezizomycotina and Saccharomycotina.** For each amino acid, its difference in the average amino acid frequency between Pezizomycotina and Saccharomycotina was calculated on the complete proteomes, which were predicted with AUGUSTUS v3.3.1 using *Aspergillus fumigatus* and *Saccharomyces cerevisiae* S288C trained models for Pezizomycotina and Saccharomycotina, respectively. Red bars denote amino acids whose average frequency is higher in Pezizomycotina than in Saccharomycotina, whereas green bars denote amino acids whose average frequency is higher in Saccharomycotina than in Pezizomycotina. This plot shows that Pezizomycotina genomes generally contain higher frequencies of amino acids encoded by GC-rich codons but lower frequencies of amino acids encoded by AT-rich codons compared to Saccharomycotina.

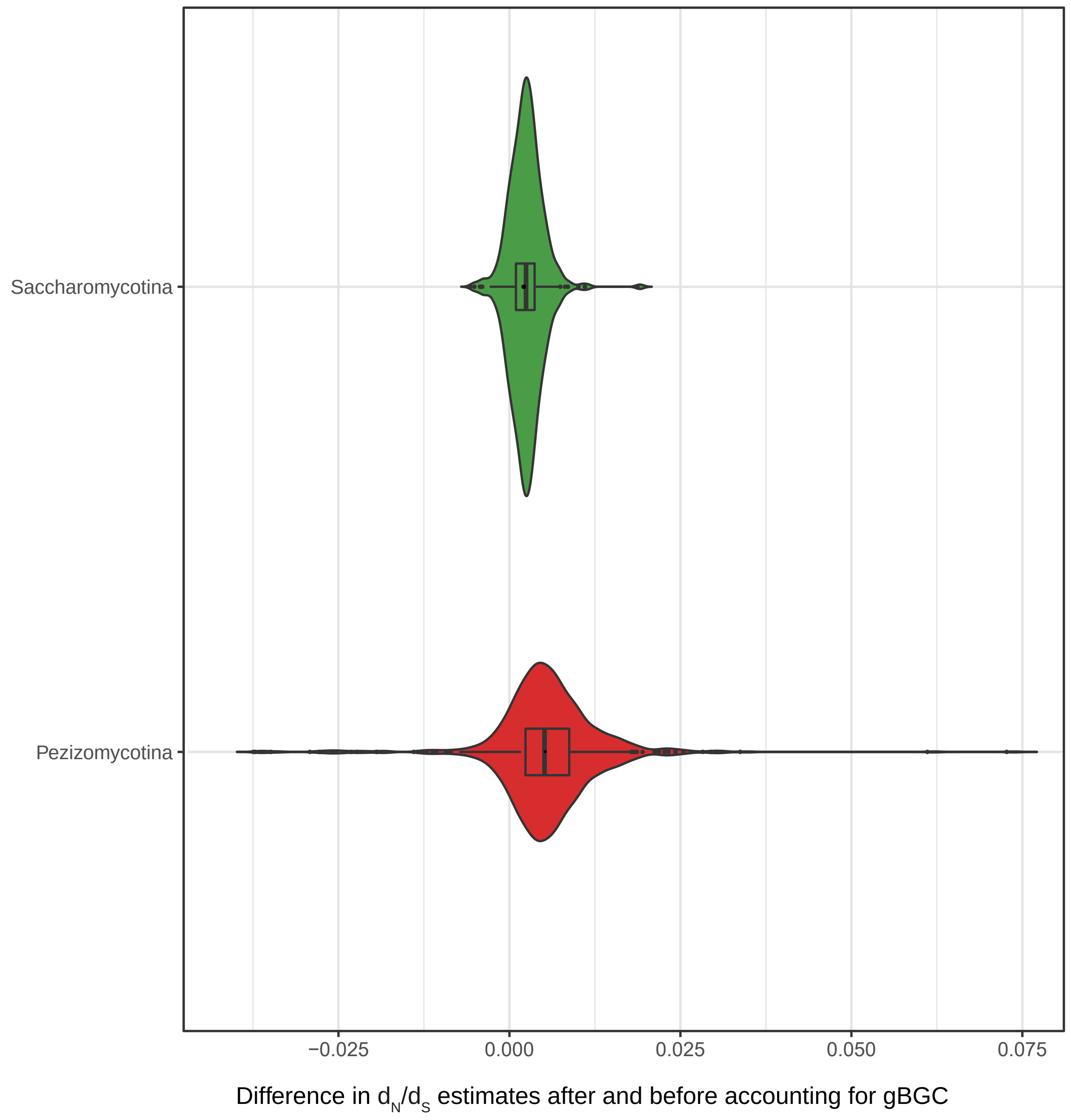


**Supplementary Figure 8. The impact of GC-biased gene conversion (gBGC) on $d_N/d_S$ estimates.** To investigate the effect of GC content on selection acting on protein-coding genes, we used an approach that accounts for non-stationary base composition, to calculate difference in $d_N/d_S$ before and after accounting for gBGC for 815 codon-based BUSCO genes in Pezizomycotina and Saccharomycotina, respectively. This plot shows that Pezizomycotina with higher GC content would suffer from greater effect of gBGC on estimating $d_N/d_S$ than Saccharomycotina with lower GC content. Calculating differences in $d_N/d_S$ after and before accounting for gBGC across 815 codon-based BUSCO genes revealed that the effect of gBGC on dN/dS estimates in Pezizomycotina is 2-fold higher than that in Saccharomycotina (Pezizomycotina: average of differences in $d_N/d_S$ = 0.004; Saccharomycotina: average of differences in $d_N/d_S$ = 0.002).
